# Midbrain organoids with an *SNCA* gene triplication display dopamine-dependent alterations in network activity

**DOI:** 10.1101/2025.04.23.650231

**Authors:** Ghislaine Deyab, Rhalena A. Thomas, Xue Er Ding, Jialun Li, Julien Sirois, Zaid Al-Azzawi, Simeng Niu, Thomas M. Durcan, Edward A. Fon

**Affiliations:** Department of Neurology and Neurosurgery, Montreal Neurological Institute-Hospital, McGill University, Montreal, Quebec H3A 2B4, Canada; The Neuro’s Early Drug Discovery Unit (EDDU), McGill University, Montreal, QC H3A 2B4, Canada

## Abstract

Human midbrain organoids (hMOs) show promise as a patient-derived model for the study of Parkinson’s disease (PD). Yet, much remains unknown about how accurately hMOs recapitulate key features of PD in the human brain. In both PD patients and animal models, disease progression leads to characteristic changes in neural activity throughout the basal ganglia. Here we demonstrate that patient-derived induced pluripotent stem cell (iPSC) hMOs harboring a triplication in the *SNCA* gene, encoding α-synuclein, a key protein in PD pathogenesis, can recapitulate PD-associated changes in neural activity. Namely, we observe hyperactive network activity in *SNCA* triplication hMOs, but not in isogenic, CRISPR-corrected iPSC hMOs. These changes are characterized by an increase in the number of bursts and network-wide bursts. Moreover, *SNCA* triplication hMOs exhibit an increase in network synchrony and burst/network burst strength similar to observations in animal and human PD brains. Subsequently, we show that the observed changes in neuronal activity are attributed to dopamine D2 receptor hypoactivity due to dopamine depletion, which could be reversed by the D2 receptor agonist quinpirole. Thus, hMOs faithfully model network wide electrophysiological changes associated with PD progression and serve as a promising tool for PD research and personalized medicine.

## Introduction

Parkinson’s disease (PD) is a neurodegenerative disorder characterized by the progressive dysfunction and loss of dopamine (DA) neurons in the substantia nigra pars compacta (SNpc), which explain many of the motor features of PD such as tremor, bradykinesia, and rigidity^1–3^. On a cellular and molecular level, neuronal dysfunction and ultimately degeneration are believed to involve lysosomal impairment, mitochondrial defects, protein misfolding, and synaptic/network dysfunction. On a neuropathological level, the formation of inclusions in neuronal cell bodies and neurites termed “Lewy bodies” (LB) and “Lewy neurites” (LN) respectively, have been recognized as one of the main hallmarks of the PD brain^4^. LBs and LNs are composed primarily of misfolded, aggregated, and phosphorylated forms of the α-synuclein (α-syn) protein^5,6^, encoded by the *SNCA* gene. The findings that point mutations (e.g. A53T) and copy-number variations (e.g. triplications) in *SNCA* lead to familial forms of PD^7,8^, followed by evidence of prion-like spreading of pathological α-syn in the nervous system^9–13^, have further consolidated the role of α-syn in PD pathogenesis. In addition to these cellular and molecular characteristics, there is a growing body of evidence that shows electrophysiological dysfunction throughout the basal ganglia and midbrain with the progression of PD^14–19^. In the basal ganglia, DA transmission plays a central and complex role in mediating neural activity by modulating downstream D1/D2 receptor expressing medium spiny neurons. Thus, progressive DA depletion through selective degeneration of nigrostriatal neurons in PD can alter downstream neuronal output and lead to aberrant network excitability.

The majority of studies examining the consequences of DA depletion on neuronal activity have been carried out in animal models^20,21^. Although able to mimic certain aspects of the circuitry and *in vivo* processes in the human brain, animal models lack the corresponding human genetic background. Furthermore, animal studies often resort to mechanical or chemical lesioning of SNpc neurons, which does not faithfully mimic the progressive degenerative process that occurs in PD. Human derived induced pluripotent stem cells (iPSCs) differentiated into DA neurons using two-dimensional (2D) cell culture systems have recently gained traction as a PD model, as they are human specific, can be derived from patients harboring disease mutations, and are readily accessible for electrophysiological recording^22–23^. However, as a three-dimensional (3D) structure, the brain provides a rich spatial framework that vastly enhances the complexity of signaling among and between various cell types. As DA neurons in 2D cell culture inherently lack this architectural complexity, they likely fail to capture much of the intricate signaling that occurs in both the healthy and diseased human brain during PD progression. Thus, better models are required.

Recently developed iPSC-derived 3D cerebral organoids show promise in bridging the gap between traditional 2D cell culture models and *in vivo* animal models by better capturing key features of the human brain at a genetic, molecular, and physiological level^24^. Characterization of these organoids has revealed the emergence of complex network activity, reflecting aspects of early brain development^25^. Moreover, advances in cerebral organoid models have allowed for the generation of region-specific organoids, including forebrain, hindbrain, and midbrain organoids^26–29^. For instance, human midbrain organoids (hMOs), which have been developed to model the brain region containing SNpc DA neurons affected in PD, have been recently used to recapitulate key aspects of PD pathology. In particular, hMOs derived from patients with PD-associated mutations were shown to develop progressive α-syn pathology and loss of DA neurons^30–33^. However, to date there have been no in-depth studies using hMO models of PD to investigate the electrophysiological deficits that have been observed in animal models of PD and PD patients. Here we used multielectrode arrays (MEA) to determine the electrophysiological network activity in patient-derived iPSC hMOs harboring a *SNCA* triplication (*SNCA* Trip) in comparison to isogenic iPSC-derived control hMOs (Iso CTL), in which the PD-linked mutation has been CRISPR-corrected to wild-type^34,35^. We demonstrate that *SNCA* Trip hMOs display aberrant activity that emerges with age (after 5 months in culture) and coincides with an increase in molecular pathology, including an increase in phosphorylated α-syn (pSyn) inclusions, and a decrease in the number of tyrosine hydroxylase (TH) expressing DA neurons. Interestingly, we also show that activation of D2 receptors in *SNCA* Trip hMOs reverses the aberrant activity.

These findings highlight a key role for D2 receptor hypoactivity in electrophysiological dysfunction in *SNCA* Trip hMOs and illustrates the potential of hMOs to study aberrant network activity in PD.

## Results

### *SNCA* Trip hMOs show key characteristics of PD pathology

We generated PD patient iPSC-derived *SNCA* Trip and Iso CTL hMOs (Fig. 1A), which we have shown previously to contain a heterogeneous population of both neuronal and glial cell types^30,34,36,37^. Here we confirm mature neurons (MAP2+), DA neurons (TH+), astrocytes (GFAP+), inhibitory GABA neurons (GAD67+), and neural precursor cells (NES+) are present in hMOs at 5 months via immunofluorescence (Fig. 1B). We further validated the presence of glial cells and different neuronal subtypes (both young and old), via flow cytometry analysis through differential expression of 9 different cell surface markers and the intracellular DA marker TH at 3 months (Fig. 1C), using our CellTypeR workflow^37^. To confirm the increased expression of α-syn protein in *SNCA* Trip hMOs, we performed immunofluorescence analysis in 5-month-old hMOs. We show an increase in total α-syn intensity in *SNCA* Trip hMOs relative to Iso CTL hMOs, but no change in the number of α-syn+ cells (Fig.1D-E), indicating that, in the SNCA Trip hMOs, α-syn is expressed at higher levels per cell rather than in more cells.

**Figure 1.**
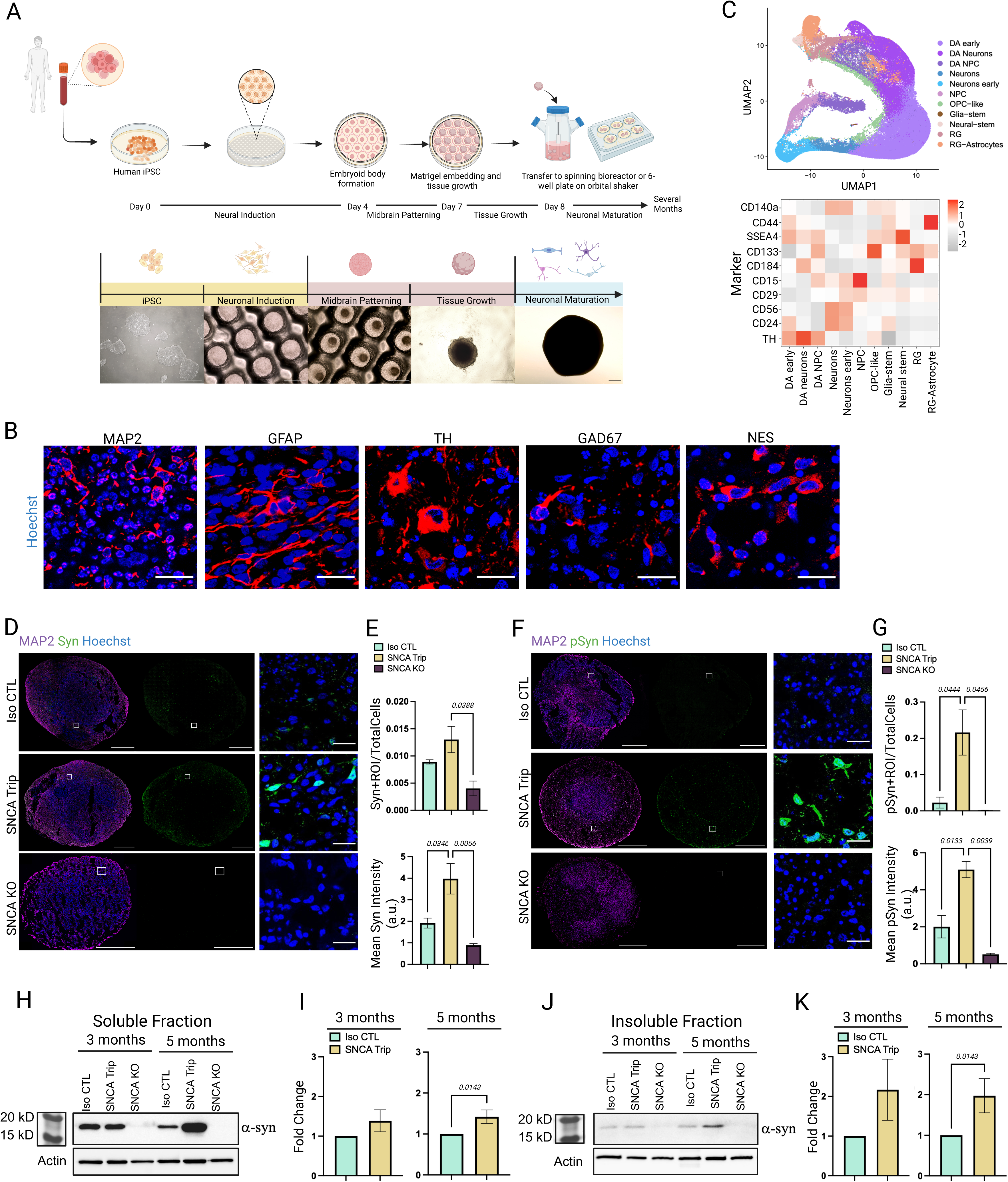
Characterization of iPSC derived *SNCA* Trip and Iso CTL hMOs. **A)** Graphical representation of hMO generation starting from iPSCs and undergoing neuronal induction, midbrain patterning, and cell maturation. Representative images are shown with bright field microscopy for each step (iPSCs in 6cm petri dish, embryoid bodies in EB DISK, and hMOs in 6-well plate). **B)** Immunofluorescence staining showing the presence of neurons (MAP2), astrocytes (GFAP), dopaminergic neurons (TH), inhibitory neurons (GAD67), and neural progenitor cells (NES) in 5-month-old hMOs. Scale bars = 25 µm. **C)** Top: A representative UMAP of the identified cell types in 3-month-old hMOs via flow cytometry. Bottom: Heatmap of the average antibody expression per identified cell cluster. (N=1) **D)** Immunofluorescence staining of total synuclein in 5-month-old Iso CTL, *SNCA* Trip, and *SNCA* KO hMOs. Scale bars on whole hMO images are 500 µm, and on close-up representative images are 50 µm. **E)** Analysis of synuclein expression shown in panel **D.** Synuclein cell count represents the total number of cells containing cytoplasmic synuclein normalized to the total number of cells detected. (N=3) **F)** Immunofluorescence staining of pSyn in 5-month-old Iso CTL, *SNCA* Trip, and *SNCA* KO hMOs. Scale bars on whole hMO images are 500 µm, and on close-up representative images are 50 µm. **G)** Analysis of pSyn expression shown in panel **F.** (N=3) **H)** Representative immunoblot against α-syn (MJFR) on the soluble protein fraction from Iso CTL, *SNCA* Trip, and *SNCA* KO hMOs at 3 and 5 months old. **I)** Quantification of soluble α-syn immunoblot at both time points. (N=3) **J)** Representative immunoblot against α-syn (MJFR) on the insoluble protein fraction from Iso CTL, *SNCA* Trip, and *SNCA* KO hMOs at 3 and 5 months old. **K)** Quantification of insoluble α-syn immunoblot at both time points. (N=3) Created in BioRender. Deyab, G. (2025) https://BioRender.com/n84x119

Moreover, hMOs generated from iPSCs derived from the same genetic background as the *SNCA* Trip hMOs, in which all copies of the *SNCA* gene were knocked-out with CRISPR (*SNCA* KO)^35^, showed only low background staining, confirming the specificity of the α-syn antibody (Fig.1D-E). Subsequently, we performed immunofluorescence analysis against pSyn, a marker of pathological α-syn aggregates in LBs and LNs^4–5^. In 5-month-old hMOs we observed an increase both in pSyn intensity and in the number of pSyn+ cells in *SNCA* Trip hMOs relative to Iso CTL hMOs (Fig.1F-G), consistent with previous findings^30^. As expected, *SNCA* KO hMOs did not show any specific pSyn staining (Fig.1F-G). Consistent with the pSyn immunofluorescence data above (Fig.1F-G), fractionation of soluble and insoluble protein from *SNCA* Trip hMOs showed significantly increased α-syn levels in both the soluble (Fig.1H-I) and insoluble (Fig.1J-K) fractions relative to Iso CTL hMOs at 5 months, but not at an earlier timepoint (3 months). These results indicate an age-dependant increase in endogenous aggregated α-syn species in the *SNCA* Trip hMOs relative to Iso CTL hMOs.

### Network-wide oscillatory activity emerges with age and is altered in *SNCA* Trip hMOs

Although previous studies have shown the emergence of spontaneous activity in hMOs^38–40^, the pattern of activity and how it changes across development is less clear. Using a 16-electrode MEA (Axion Biosystems), to assess basal activity across development, we recorded the total number of neuronal local field potentials (referred to as spikes), bursts, and network-wide bursts (periods where bursting occurs across multiple electrodes in synchrony) in Iso CTL hMOs cultured for 1, 3, and 5 months (Fig.2A). We observed a prominent increase in activity over time in culture, marked by an increase in spikes, bursts, and network bursts (Fig.2B). We find that prominent bursting and network bursting emerge between 3-5 months of age (Fig.2B-C). The emergence of bursting activity reflects neuronal maturation, whereas the occurrence of network bursting activity implies an increase in connectivity between neurons and the emergence of neural networks. Activity recordings were validated for MEA artifacts with the addition of 3uM tetrodotoxin (TTX), a voltage gated sodium channel inhibitor. The incubation of hMOs with TTX treatment led to an abolishment of neuronal activity (Fig.2D-E).

**Figure 2.**
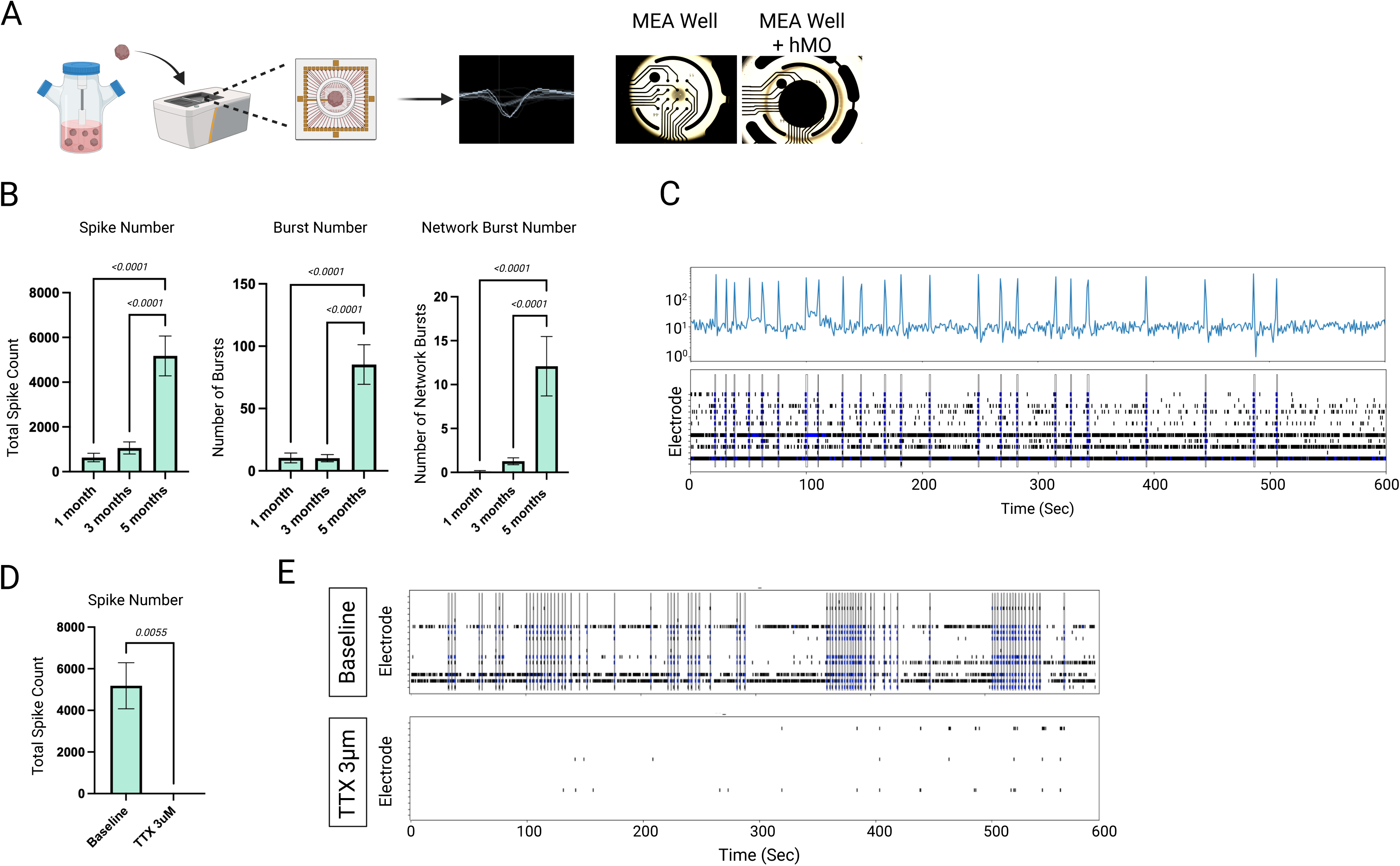
Characterization of spontaneous neuronal activity across hMO development. **A)** Graphical schematic showing MEA workflow and a representative image taken from Axion Biosystems MEA showing a local field potential waveform from a single electrode. Brightfield images showing the 16 electrodes of a single MEA well without and with an hMO on top. **B)** Quantification of neuronal activity development in Iso CTL hMOs, taken from a 10-minute MEA recording. (N=4) **C)** Representative raster plot from a 5-month-old Iso CTL hMO showing the presence of neuronal bursts (spikes highlighted in blue) and network bursts (vertical boxes). The raster plot is accompanied by spike histogram showing the summation of spikes across all electrodes in 1 second bins. **D)** Quantification of TTX treatment on Iso CTL hMOs taken from a 10-minute MEA recording. Total number of spikes is compared before and after treatment with 3µm TTX. (N=1) **E)** Representative raster plots of a single hMO before and after treatment with TTX showing the abolishment of neuronal activity. Created in BioRender. Deyab, G. (2025) https://BioRender.com/i13b607

Studies have shown shifts in the organization of neuronal activity with progressive depletion of DA in PD, primarily characterized by an increase in network synchrony and burst strength^17,41,42^. To assess whether aberrant activity also emerges in *SNCA* Trip hMOs, we compared patterns of neuronal activity between *SNCA* Trip and CTL hMOs at three timepoints (1,3, and 5 months). We see no change in the total number of spikes detected by the MEA between *SNCA* Trip and CTL hMOs at any time points (Fig.3A). Conversely, we did observe an increase in burst number and network burst number at 5 months in the *SNCA* Trip hMOs relative to Iso CTLs (Fig.3B-C), indicating that the observed differences in burst activity are not due to individual neurons firing more often, but due to a shift in their firing pattern. Therefore, we investigated how patterns of neuronal activity differed between 5-month-old *SNCA* Trip and Iso CTL hMOs. Burst and network burst strength are defined by the average number of spikes present in bursts/network bursts, as well as average burst/network burst duration. *SNCA* Trip hMOs displayed greater burst strength, as shown by longer bursting periods with more spikes per burst (Fig. 3B), longer network burst periods with more spikes per network burst (Fig. 3C, E), and increased network synchrony compared to Iso CTL hMOs (Fig. 3D, E). This shift in bursting strength can be observed by a shift in inter spike interval (ISI) distribution between the Iso CTL and the *SNCA* Trip hMOs (Fig. 3F). The ISI distribution of a mature neural network is bimodal, with a peak of short ISIs (representing fast spiking within bursting periods), and a peak of long ISIs (representing sparse spiking between bursting periods). Here, we observe a shift in the ISI distribution of *SNCA* Trip hMOs at 5-months-old towards having a larger peak of short ISIs and a smaller peak of long ISIs (Fig. 3F). This shift is representative of a higher frequency of spikes during burst periods and a lower frequency of spiking outside of bursting periods (Fig. 3G). No change in the ISI of spikes within bursting periods is observed between genotypes (Fig. 3H), suggesting that neurons within bursting periods are not firing faster. Thus, the increased number of spikes within a burst could be accounted for by the increased burst duration. To further explore the contributions of the different network burst features to the patterns of electrophysiological activity observed within *SNCA* Trip and Iso CTL hMOs, we performed a principal component analysis (PCA), a dimensionality reduction method that permits visualization of multiple parameters. Plotting the first 2 principal component values for each hMO, we observed that *SNCA* Trip hMOs cluster separately from Iso CTL hMOs (Fig. 3I). We further visualized the correlation between our input parameters and the principal components to see which parameters most describes the clustering of our hMOs in latent space. We observed that *SNCA* Trip hMOs cluster towards the quadrant more strongly described by parameters representing burst and network burst strength (Fig. 3J). Additionally, independent hierarchical clustering of *SNCA* Trip and Iso CTL hMOs based on variables used to describe burst strength (burst duration, spikes per burst, network burst duration, spikes per network burst) generally show separate clustering of the two genotypes (Fig. 3K). This highlights that the differences in firing patterns between the two genotypes are primarily based on their burst/network burst strength. Taken together, these data indicate that *SNCA* Trip hMOs display a shift in their neuronal firing patterns to produce stronger bursting and network bursting periods and increased neural network synchrony. This shift in *SNCA* Trip hMO neuronal activity emerges between 3 and 5 months, coinciding with the emergence of increased pSyn expression and pathological α-syn aggregates (Fig.1F-K). The emergence of both α-syn pathology and aberrant network activity at this timepoint raises the question of whether the increase in α-syn levels, aggregation, and phosphorylation impact network activity directly or indirectly through the downstream consequences of neurodegeneration and DA depletion.

**Figure 3.**
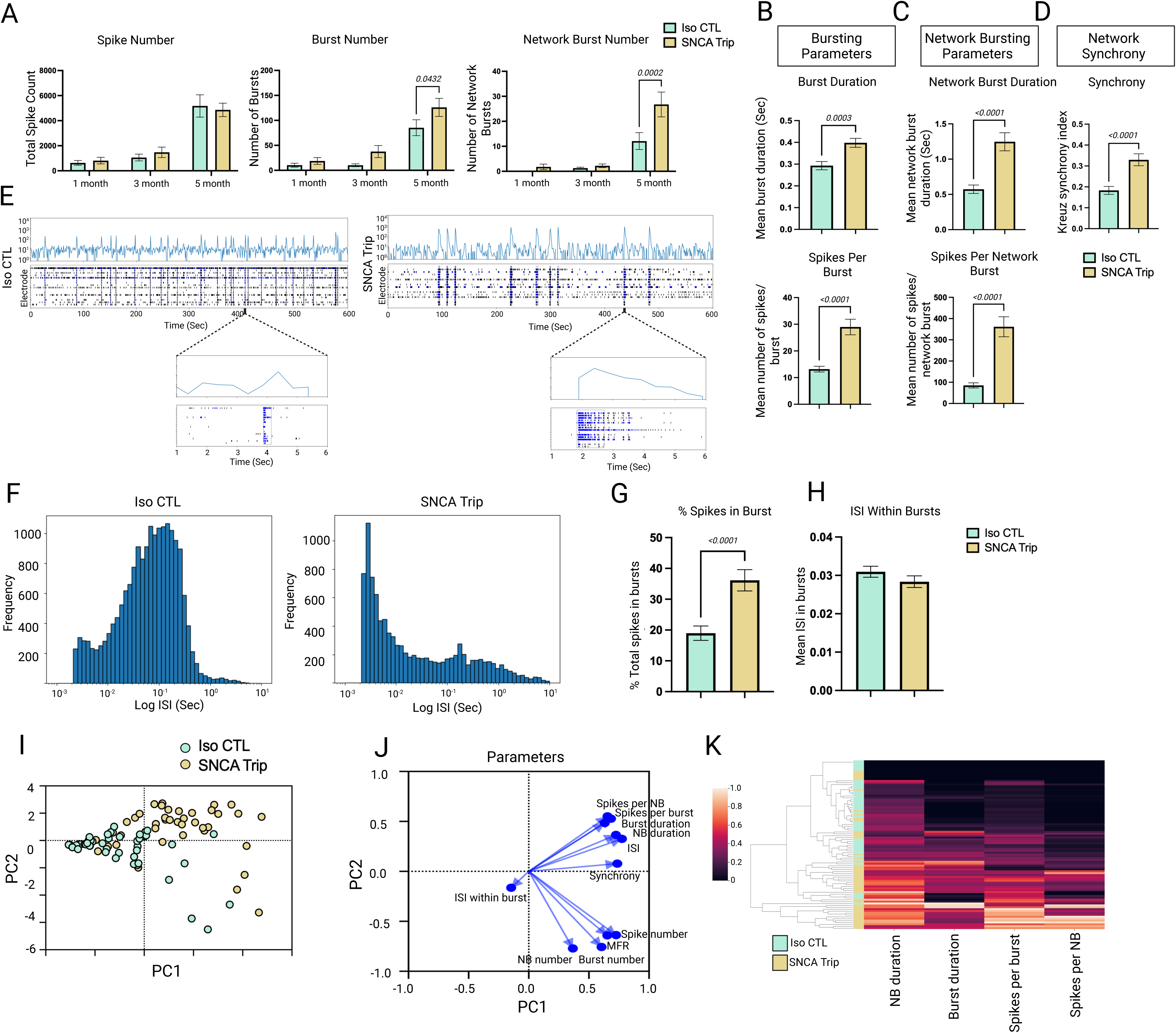
Comparison of spontaneous neuronal activity between *SNCA* Trip and Iso CTL hMOs. **A)** Quantification of neuronal activity between Iso CTL and *SNCA* Trip hMOs taken from 10-minute MEA recordings across development. Differences are observed at 5-months-old in the number of bursts and network bursts produced but not in the number of total spikes between the two genotypes. (N=4) **B)** Quantification of bursting parameters between Iso CTL and *SNCA* Trip hMOs taken from a 10-minute MEA recording at 5-months-old. (N=4) **C)** Quantification of network bursting parameters between Iso CTL and *SNCA* Trip hMOs taken from a 10-minute MEA recording at 5-months-old. (N=4) **D)** Quantification of synchrony between Iso CTL and *SNCA* Trip hMOs taken from a 10-minute MEA recording at 5-months. (N=4) **E)** Representative raster plots of a single Iso CTL and *SNCA* Trip hMO at 5-months-old (showing neuronal bursts highlighted in blue and network bursts outlined by a vertical black box) accompanied by a spike histogram above showing the summation of spikes across electrodes per 1 second bin. Zoomed in raster plots show a 5 second interval containing a network burst to highlight the difference in network burst strength between Iso CTL and *SNCA* Trip hMOs. **F)** Representative ISI distribution of a single Iso CTL and *SNCA* Trip hMO from a 10-minute MEA recording showing a shift towards larger ISIs in Iso CTL hMOs compared to *SNCA* Trip indicating fewer spikes involved in bursting periods. **G)** Quantification of the average percent of total spikes involved in bursts between Iso CTL and *SNCA* Trip hMOs at 5-months-old. (N=4) **H)** Quantification of the average ISI within bursting periods between Iso CTL and *SNCA* Trip hMOs at 5-months-old. (N=4) **I)** PCA plot showing the separate clustering of Iso CTL and *SNCA* Trip hMOs at 5-months-old based on the parameters from panel **J.** Separation of clusters is primarily based on parameters describing burst and network burst strength. (N=2) **J)** The correlation of the measured parameters (spike number, burst number, network burst number, MFR, ISI, ISI in bursts, burst duration, network burst duration, spikes per burst, spikes per network burst, Kreuz synchrony index) to the principal components used for PCA between *SNCA* Trip and Iso CTL hMOs at 5-months-old. Parameters that group in a given quadrant more strongly represent hMOs that cluster in that same quadrant. (N=2) **K)** Independent hierarchical clustering of Iso CTL and *SNCA* Trip hMOs showing *SNCA* Trip hMOs have stronger burst/network burst durations and an increased number of spikes per burst/network burst. (N=2) Created in BioRender. Deyab, G. (2025) https://BioRender.com/b39r941

### D2 receptor activation rescues aberrant activity in *SNCA* Trip hMOs

To investigate the role of DA in modulating patterns of network activity, we first determined the levels of TH, the rate-limiting enzyme in DA biosynthesis, in *SNCA* Trip and Iso CTL hMOs using immunoblot analysis. Whereas no significant differences were observed between genotypes in 3-month-old hMOs, we find a significant decrease in TH levels in *SNCA* Trip hMOs at 5 months, supporting a time-dependent loss rather than a developmental defect in TH expression (Fig.4A-B). We further validated the loss of TH expressing neurons via immunofluorescence and again observe a significant decrease in the number of TH positive cells in *SNCA* Trip hMOs relative to Iso CTLs in 5-month-old hMOs (Fig.4C-D), confirming our previous findings using flow cytometry^30^. Together, these findings indicate a decrease in DA biosynthetic capacity and number of DA neurons in *SNCA* trip hMOs at the age which correlates with the emergence of pSyn neuropathology and aberrant network activity.

**Figure 4.**
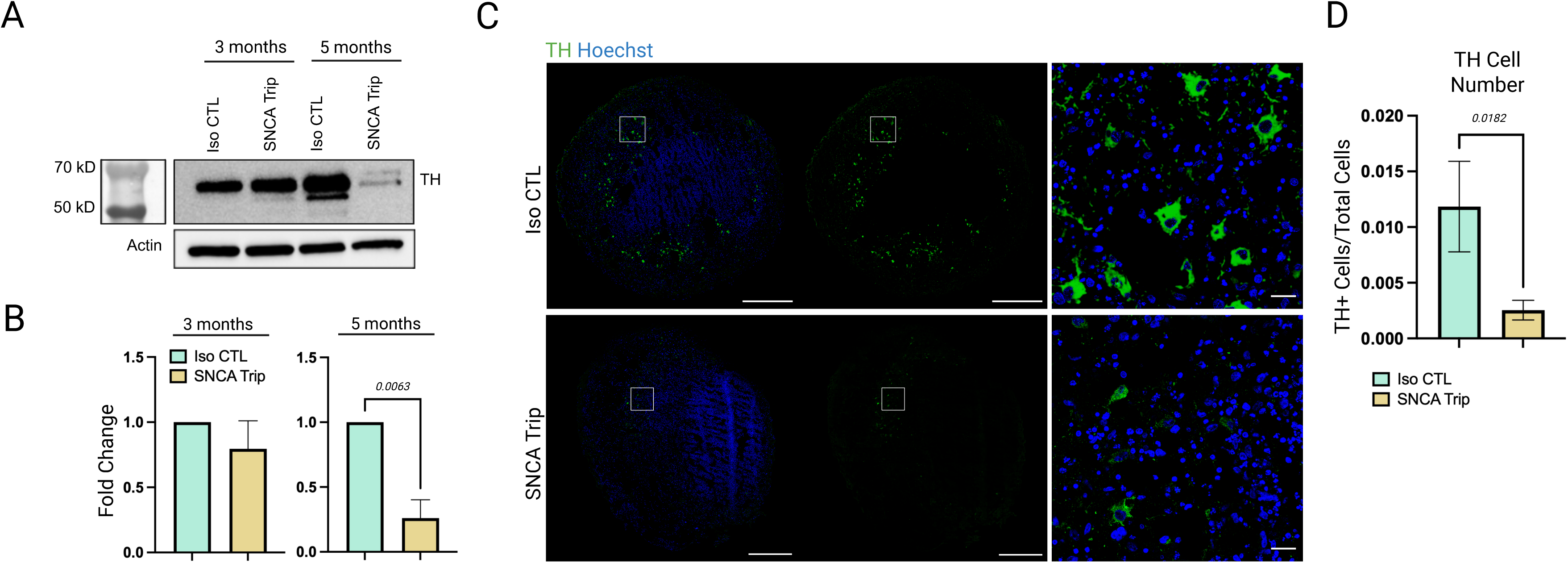
Comparison of DA neurons in *SNCA* Trip and Iso CTL hMOs. **A)** Immunoblot analysis on TH protein levels in Iso CTL and *SNCA* Trip hMOs taken at 3 and 5 months, showing a decrease in TH expression in *SNCA* Trip hMOs at 5-months-old but not earlier. **B)** Quantitative analysis of immunoblot shown in panel **A**. (N=3) **C)** Immunofluorescence staining against TH expression in Iso CTL and *SNCA* Trip hMOs at 5-months-old. Scale bars on whole hMO images are 500 µm and on zoomed in representative images are 25 µm. **D)** Quantification of TH positive cells in panel **C.** (N=3) Created in BioRender. Deyab, G. (2025) https://BioRender.com/d28l622

One explanation for the aberrant activity in *SNCA* Trip hMOs is that reduced DA production leads to changes on DA receptor activation. In particular, D2 receptors which are expressed both pre-synaptically on DA neurons and post-synaptically on medium spiny GABA neurons, have been previously identified as key players in mediating aberrant basal ganglia activity in cellular and mouse models of PD^40,42^. Inhibitory D2 receptors induce hyperpolarization of the cell membrane in response to DA transmission. It is therefore thought that a decrease in DA transmission in PD leads to decreased inhibition of downstream D2 receptor-expressing neurons, leading to a hyperactive basal ganglia circuitry. Therefore, to test if D2 receptor activity is affected by the observed decrease in DA biosynthetic capacity and DA neuron numbers in *SNCA* Trip hMOs, we treated 5-month-old hMOs with quinpirole (5uM for 15 minutes), a selective D2/D3 receptor agonist before recording network activity. Quinpirole significantly reduced the number of spikes, bursts, and network bursts in Iso CTL hMOs (Fig. 5A), pointing to the inhibitory nature of D2 receptor activation in wild-type conditions. In contrast, in *SNCA* Trip hMOs, quinpirole treatment significantly reduced network burst number, but not the total number of spikes or bursts (Fig.5A). These results indicate that, in *SNCA* Trip hMOs, quinpirole counteracts the reduction in D2 receptor activation, rescuing the observed network burst hyperactivity down to levels comparable to untreated Iso CTLs hMOs but with little effect on individual neuron activity, as manifested by the unaltered spikes and bursts. We next wanted to determine the effects of D2 receptor activity on the organization of spikes within bursting and network bursting periods, as shown above (Fig. 3B-F). Quinpirole treatment rescued both burst (Fig. 5B) and network burst (Fig. 5C) strength in *SNCA* Trip hMOs, as measured by the duration and the number of spikes per burst/network burst, to levels comparable to Iso CTL hMOs. Similarly, quinpirole treatment significantly reduced network synchrony (Fig. 5D). This rescue of network activity toward control levels is also evident in the raster plot visualization (Fig. 5E) and in the rescue of the ISI distributions of *SNCA* Trip hMOs after quinpirole treatment to those seen in Iso CTL hMOs (Fig.5F). We see no change in burst duration in Iso CTL hMOs with and without quinpirole treatment (Fig.5B,E), however we observe decreased overall burst production in Iso CTL hMOs treated with quinpirole which is also consistent with the shift in their ISI distribution to more spikes with longer ISIs (Fig.5F).

**Figure 5.**
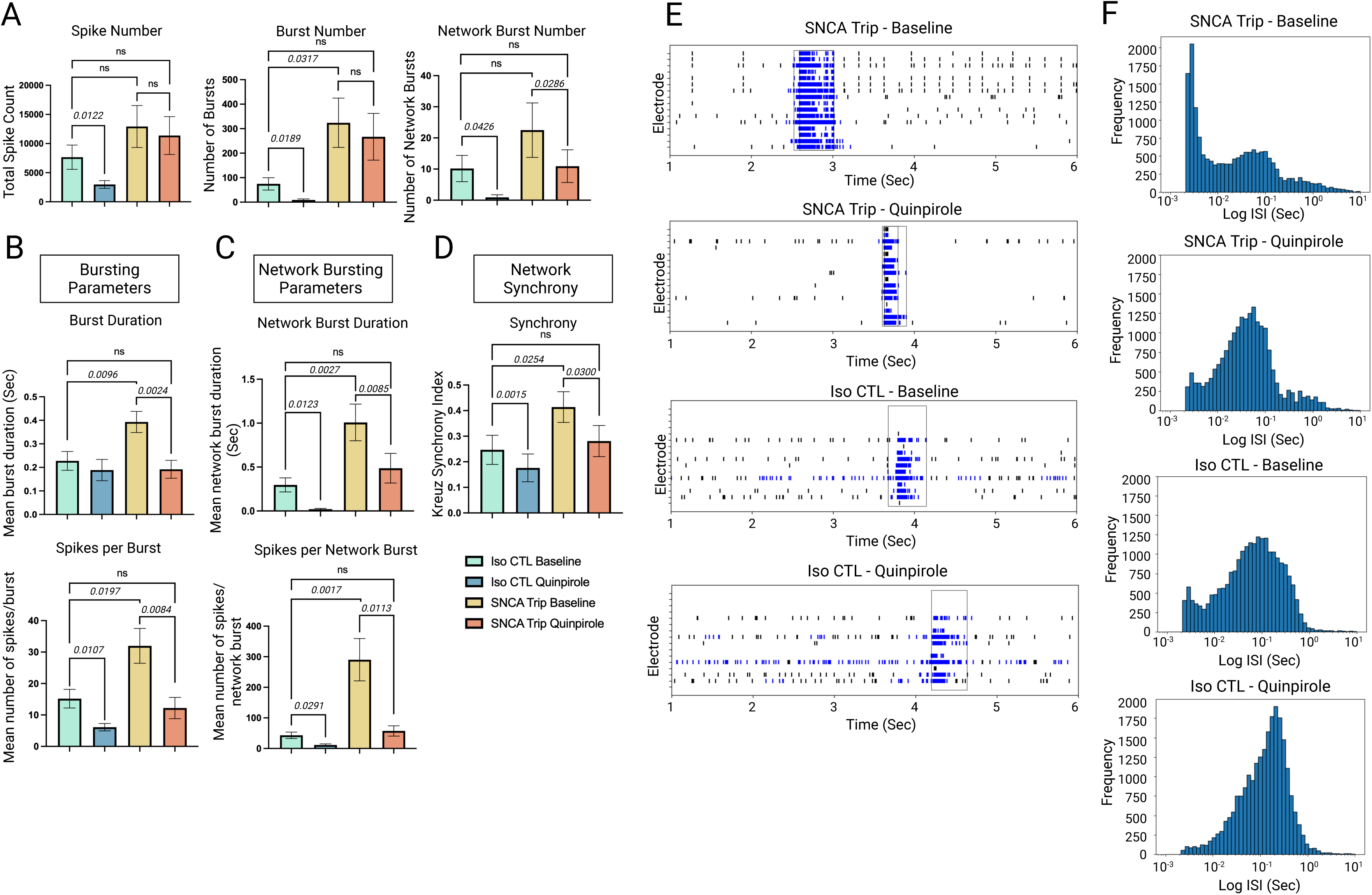
Assessing the role of D2 receptors in modulating patterns of hMO neuronal activity. **A)** Quantification of quinpirole treatment on 5-month-old Iso CTL and *SNCA* Trip hMOs activity taken from 10-minute MEA recordings. (N=3) **B)** Quantification of quinpirole treatment on Iso CTL and *SNCA* Trip hMO bursting parameters. (N=3) **C)** Quantification of quinpirole treatment on network bursting parameters. (N=3) **D)** Quantification of quinpirole treatment on synchrony. (N=3) **E)** Raster plot representation of a network burst before and after quinpirole treatment in a Iso CTL and *SNCA* Trip hMO at 5-months-old. Recordings were taken from the same hMO before and after 15 minutes of quinpirole treatment. A 5 second interval of each recording is shown in the raster plot. **F)** Representative ISI distribution taken from 10-minute MEA recordings from a Iso CTL and *SNCA* Trip hMO at 5-months-old before and after treatment with quinpirole showing a shift in ISI distribution to larger ISIs in quinpirole treated hMOs, resembling Iso CTL hMOs at a matched age. Created in BioRender. Deyab, G. (2025) https://BioRender.com/c01s398

Together, these observations imply that *SNCA* Trip hMOs display D2 receptor hypoactivity that contributes to downstream neuronal network hyperactivity, which can be rescued by D2 receptor activation.

### DA depletion in Iso CTL hMOs mimics network hyperactivity seen in *SNCA* Trip hMOs

To test whether reducing DA transmission in wild-type hMOs is sufficient to recapitulate the aberrant network activity observed in PD patient-derived *SNCA* Trip hMOs, we depleted DA stores by chronically treating Iso CTL hMOs with tetrabenazine (TBZ), a high affinity inhibitor of VMAT2. TBZ treatment was started at 3-months in culture, continued for 2 consecutive months, and recordings were performed in 5-month-old hMOs. This was done to mimic the period during which PD pathology develops and a decrease in TH expressing DA neurons is observed in *SNCA* Trip hMOs. After 2 months of TBZ treatment, Iso CTL hMOs exhibited a significant increase in burst (Fig.6A) and network burst (Fig.6B) strength compared to untreated hMOs, phenocopying the aberrant network activity observed in *SNCA* Trip hMOs (Fig.3A-B).

**Figure 6.**
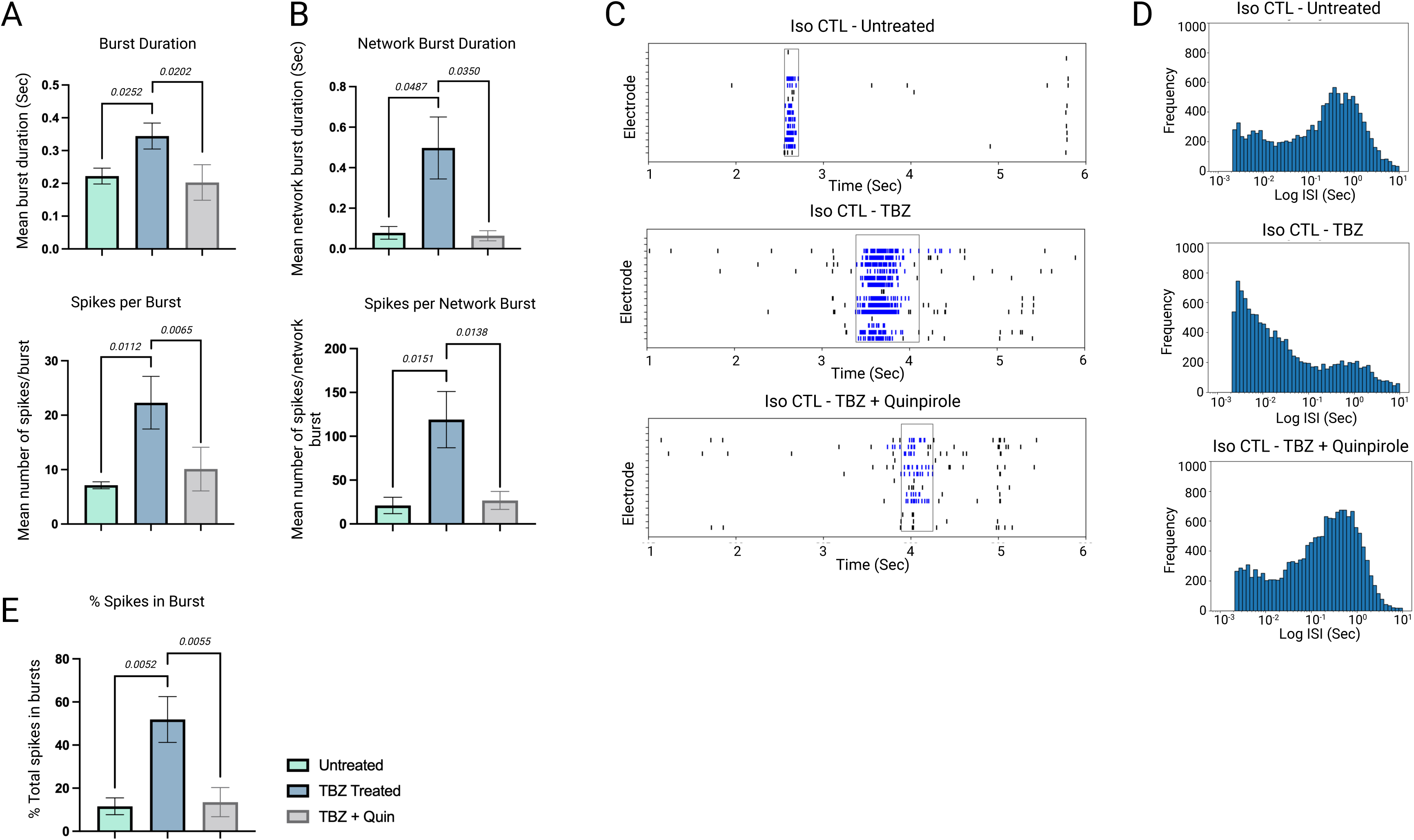
Assessing the role of DA depletion in modulating patterns of hMO neuronal activity. **A)** Quantification of bursting parameters in Iso CTL hMOs taken from a 10-minute MEA recording either untreated, chronically treated with TBZ (TBZ treated), or chronically treated with TBZ followed by subsequent treatment with quinpirole 15 minutes prior to recording (TBZ + Quin). (N=1) **B)** Quantification of network bursting parameters in Iso CTL hMOs either untreated, TBZ treated, or TBZ + Quin Treated. (N=1) **C)** A 5 second raster plot representation of a network burst recorded from an untreated Iso CTL hMO, a TBZ treated Isp CTL hMO, and a TBZ + Quin treated Iso CTL hMO at 5-months-old. **D)** Representative ISI distributions of a untreated, TBZ treated, and TBZ + Quin treated Iso CTL hMO at 5-months-old. **E)** Quantification of the percent of total spikes involved in bursting periods between untreated, TBZ treated, and TBZ + Quin treated hMOs. (N=1) Created in BioRender. Deyab, G. (2025) https://BioRender.com/r06z537

Subsequent treatment with quinpirole rescued the TBZ-induced increase in burst/network burst strength to levels comparable to untreated Iso CTL hMOs (Fig.6A-B), similar to what was observed in *SNCA* Trip hMOs (Fig.4B-C). This is further illustrated by raster plot visualization and ISI distribution in which TBZ treated Iso CTL hMOs display longer network bursts with a shift in their ISI distribution to an increased abundance of spikes with short ISIs, mimicking what was observed in SNCA Trip hMOs (Fig.6C-D). Again, these changes could be reversed by quinpirole (Fig.6C-D). This is further validated by the observed increase in the percent of total spikes within bursts in TBZ-treated Iso CTL hMOs, which is rescued by quinpirole treatment (Fig.6E), again phenocopying findings in *SNCA* Trip hMOs (Fig.3G). Taken together, these results indicate that the aberrant network activity associated with PD can be phenocopied in wild-type, Iso CTL hMOs by mimicking the reduction in DA transmission that occurs spontaneously in *SNCA* Trip hMOs. Moreover, the effects of reduced DA transmission on network hyperactivity appear to be mediated by D2 receptor hypoactivation, as they could be reversed by quinpirole in both PD-associated *SNCA* Trip hMOs and DA depleted wild-type Iso CTL hMOs.

## Discussion

The need for disease relevant patient-specific models in PD research is warranted. With the advent of iPSC technology, researchers have turned to more sophisticated iPSC systems such as brain organoids to more faithfully model the complexity of the brain, to better understand neurological disease pathogenesis, and to accelerate drug discovery. Several studies have reported PD pathology in hMOs^30,32,33,40,45^. Yet, given the extensive literature showing that PD leads to aberrant activity throughout the basal ganglia^15,17,42,46^, to our knowledge, none have focused on characterizing changes in network activity in hMO models of PD. Indeed, changes in network activity in PD have been proposed to play a role in prion-like propagation of α-syn via activity-dependent synaptic vesicle release^47–50^ and numerous studies have indicated that abnormal network activity is a marker of disease progression and pathology^17,18,50–53^. Here we characterize patterns of neural network activity in hMOs derived from iPSCs harboring an *SNCA* gene triplication to investigate how aberrant network activity develops in a hMO model of PD. We show that *SNCA* Trip hMOs produce shifts in neural network activity characterized by hyperactive bursting and network bursting activity, as well as an increase in synchrony across the neural network, both reminiscent of changes observed in *in vivo* animal models^17,18,42^. We further show that the increase in burst and network burst strength observed in *SNCA* Trip hMOs is a consequence of DA depletion and is dependent on D2 receptor activity. This is evident from our results showing that activating D2 receptors rescue the bursting phenotype seen in *SNCA* Trip hMOs, and chronically depleting DA stores in Iso CTL hMOs phenocopies the changes in network activity observed in *SNCA* Trip hMOs.

Experiments in rodent models of PD show increased firing rates and irregularities of neural activity throughout the basal ganglia, as well as an increased frequency of neural bursts and synchronization between basal ganglia and midbrain nuclei^15,17,42^. Willard and colleagues showed increased bursting at early stages of DA depletion but that aberrations in network wide synchrony only appear with severe DA depletion (5% TH remaining), leading to motor dysfunction. In line with the literature, we see an increase in bursting/network bursting strength and synchrony coinciding with loss of TH positive cells and pSyn accumulation at 5 months of age. In contrast, we do not see changes in network activity in *SNCA* Trip hMOs at 3 months of age, when there is no detectable loss of TH neurons or accumulation of insoluble α-syn species. We do not know if the lack of changes in network activity prior to 5 months is due to an immature neuronal network that prevents prominent network bursting activity from being captured, or the lack of PD pathology at this stage. However, we show that the network changes observed at 5 months of age are likely a response to decreased DA transmission, as depleting DA in healthy Iso CTL hMOs, without the induction of other PD pathology, mimics the observed changes in network activity. This raises questions about the role of α-syn in aberrant network activity in PD. Given α-syn’s function at the synapse, it has been proposed that pathological α-syn may contribute to electrophysiological pathology in PD^53–56^. These studies point to pathogenic α-syn reducing DA transmission and reuptake. However, here we show that reducing DA transmission in healthy organoids can mimic the electrophysiological changes associated with PD pathology. Furthermore, other genes implicated in PD that do not consistently lead to α-syn pathology also cause aberrant neuronal activity, including increased network synchrony^57,58^. Therefore, although α-syn may play a role in early neuronal dysfunction, it is likely that large disruptions in network activity leading to motor dysfunction are caused by reduced DA transmission. However, the role of pathological α-syn in synaptic dysfunction is still not completely understood and may be contributing to our observations in *SNCA* Trip hMOs. Similarly, whether synaptic dysfunction contributes to the loss of TH expressing neurons, is a compensatory mechanism following DA depletion, or a combination of the two is still unknown^59^.

Our results indicate that the observed aberrant network activity is attributed to a decrease in D2 receptor activity in *SNCA* Trip hMOs. We show that pharmacological activation of D2 receptors rescues the aberrant activity in *SNCA* Trip hMOs and DA depleted Iso CTL hMOs. D2 receptors are expressed both pre- and post-synaptically and activate G-protein activated inwardly rectifying potassium channels to hyperpolarize the cell membrane. In parallel, D2 receptor activation regulates DA release by inhibiting DA transmission, increasing dopamine transporter function, and inhibiting TH-mediated DA synthesis^60,61^. The modulatory activity of D2 receptors is dependent on extracellular DA concentrations, therefore D2 receptors function as DA sensors which can trigger feedback responses to regulate DA neuron activity^62^. The role of D2 receptor hypoactivity in emergent electrophysiological changes throughout the basal ganglia and midbrain nuclei with PD progression has been described before^41,43,63^. It is hypothesized that D2 receptor hypoactivity may be a compensatory mechanism allowing for increased DA release during early stages of PD. However, studies have shown that quinpirole treatment can prevent markers of neuronal stress and degeneration including loss of TH, reduction in network arborization, and mitochondrial damage^43,64^. Therefore, although D2 receptor hypoactivity could act as a compensatory mechanism, its hypoactivity may trigger downstream cascades that lead to increased neuronal death, such as excitotoxicity or other downstream pathways^64^.

Our results point to D2 receptor activation as a potential therapeutic for the dysfunctional network activity seen in PD. However, additional experiments are needed to determine its effects on preventing PD pathology and neurodegeneration, as proposed previously^43^. Given the rescue of aberrant network activity in both wild-type DA-depleted hMOs and in PD patient-derived *SNCA* Trip hMOs by quinpirole, it is likely that D2 receptor hypoactivation plays a prominent role in this process. Nevertheless, we cannot exclude that other receptors, ion channels, or proteins may contribute to the observed network hyperactivity in PD models, as suggested previously in mouse models^46,65–69^. However, given the robust capacity of DA depletion to induce, and D2 receptor stimulation to reverse, the aberrant network activity, we predict that these additional changes would come into play downstream of D2 receptor hypoactivity.

Importantly, our experiments were done at a timepoint when PD pathology (pSyn and insoluble α-syn accumulation, and TH cell loss) could be readily observed in *SNCA* Trip hMOs. Thus, our system is somewhat unique in mechanistically linking a human patient-derived α-syn model, which spontaneously develops α-syn pathology, reduction in DA transmission and D2 receptor hypoactivation to the electrophysiological network abnormalities shown previously in cellular or animal models of PD. Moreover, the fact that network activity emerges, and that PD-associated aberrant network activity can be modeled in hMOs is particularly remarkable given that while hMOs develop a diversity of cell types similar to the human midbrain, they do not develop the precise neural circuitry present in the intact midbrain and downstream basal ganglia nuclei. We can only conclude that the underlying cell-type diversity, neurochemistry, and signaling are sufficient to reconstitute key features of these processes, even in the absence of the appropriate anatomical circuitry. Taken together, we conclude that such hMO models could serve as a robust system to better understand the pathways and dissect the pharmacology involved in the aberrant activity and neuropathology of PD, with the aim of developing more effective and reliable medications for the disease.

## Methods

### iPSC maintenance and hMO generation

iPSCs were thawed in a 37°C water bath and plated on a Matrigel coated dish in mTeSR media and supplemented with 10µM of ROCK inhibitor (Y-27632) to promote cell survival. Media was changed daily without further addition of ROCK inhibitor, and cells were checked for spontaneous differentiation and/or contamination. The cells were passaged once they reached 60-70% confluency, a minimum of 2 passages were done before hMO generation. hMOs were generated using the protocol published by Mohamed and colleagues^34,36^. Briefly, iPSCs were washed with DMEM containing 1X Antibiotic-Antimycotic (DMEM+Anti) and enzymatically dissociated into a single cell solution with Accutase at 37°C. Accutase treatment was stopped after 5-7 minutes with the addition of DMEM+Anti, cells were collected in falcon tubes and pelleted for 3 minutes at 1200rpm. Pellets were resuspended in 1ml neuronal induction media (1:1 DMEM+Anti:Neurobasal, 1:100 N2, 1:50 B27 without vitamin A, 1% Glutamax, 1% MEM-NEAA, 5 µM 2-mercaptoethanol, 1 µg/mL Heparin, 10 µM SB431542, 200ng/mL Noggin, 0.8 µM CHIR99021, 10 µM Y-27632), and cell concentration was counted. 10,000 cells were seeded per well in a 360-well ULA-coated EB DISK (eNUVIO). EB DISKs containing iPSCs were spun down at 1200 rpm for 10 minutes and supplemented with 5mL of additional neuronal induction media and placed at 37°C with 5% CO2. After 2 days, EBs formed, and their media was changed with 5mL of neuronal induction media without ROCK inhibitor. After an additional 2 days, media was changed to midbrain patterning media (1:1 DMEM+Anti:Neurobasal, 1:100 N2, 1:50 B27 without vitamin A, 1% Glutamax, 1% MEM-NEAA, 5 µM 2-mercaptoethanol, 1 µg/mL Heparin, 10 µM SB431542, 200 ng/mL Noggin, 0.8 µM CHIR99021, 100 ng/mL SHH, 100 ng/ml FGF-8) and placed back in the incubator for 3 days. At this time, the media was removed and EBs were embedded with 1mL/EB DISK of Matrigel with reduced growth factors. The Matrigel was allowed to solidify for 30 minutes in the incubator before the addition of 5mL of tissue growth media (Neurobasal, 1:100 N2, 1:50 B27 without vitamin A, 1% Glutamax, 1% MEM-NEAA, 5 µM 2-mercaptoethanol, 200 ng/mL laminin, 2.5 µg/mL insulin, 100 ng/mL SHH, 100 ng/ml FGF-8, 1X Penicillin-Streptomycin).

EB DISCs were incubated for an additional day before they were transferred into ultra-low attachment 6-well plates or bioreactors containing final differentiation media (Neurobasal, 1:100 N2, 1:50 B27 without vitamin A, 1% Glutamax, 1% MEM-NEAA, 5 µM 2-mercaptoethanol, 10 ng/mL BDNF, 10 ng/mL GDNF, 100 µM ascorbic acid, 125 µM db-cAMP, 1X Penicillin-Streptomycin). The decision to use bioreactors or 6-well plates was based on the number of organoids generated, large batches were generated in bioreactors whereas small batches were generated in 6-well plates. All 6 well plates were placed on an orbital shaker set at 70 rpm, bioreactors were placed on a magnetic stirrer, in the incubator for the duration of their culture and an 80% media changes was performed 3 times a week. All reagents are listed in **supplementary table 1.**

### Cryosectioning and immunofluorescence staining of hMOs

hMOs were fixed in 4% PFA in PBS at 4°C overnight and then placed in a 20% sucrose-PBS solution at 4°C until hMOs were no longer floating (maximum 3 days). hMOs were then embedded in cryomolds with OCT and kept at -80°C until use. Embedded hMOs were sectioned at 10 µM thickness using a cryostat (ThermoFisher, CryoStar NX70 Cryostat), slides were stored at -20°C until further use. For immunofluorescence staining, slides were rehydrated in PBS for 15 mins before blocking (PBS, 5% NDS, 0.005% BSA, 0.2% Triton-X 100) for 1 hour at room temperature. hMO sections were then incubated in primary antibodies (**Supplementary table 2**) diluted in blocking buffer at 4°C overnight, followed by 3 x 5-minute washes in PBS and incubation with secondary antibody (**Supplementary table 2**) diluted in blocking buffer for 1 hour at room temperature. Following secondary antibody incubation, sections were washed 3 x 5-minutes in PBS and incubated in 1:5000 Hoechst diluted in PBS for 10 minutes at room temperature, followed by an additional 2 x 5-minute washes in PBS. hMO sections were then sealed with glass coverslips and AquaMount solution and subsequently imaged at 63X using the Leica TCS SP8 Confocal microscope. All confocal images were analyzed on ImageJ for staining intensity, area, and number of detected ROIs using custom analysis scripts written in python and run in Fiji (https://github.com/neuroeddu/OrgQ).

### Preparation of soluble and insoluble cell lysates from hMOs

hMOs were snap-frozen in liquid nitrogen and stored at -80°C until fractionation. For soluble/insoluble protein fractionation we followed the protocol outlined in Stojkovska & Mazzulli^70^ with some minor changes optimized for hMO tissue. Briefly, 3 frozen hMOs per condition were pooled and lysed in 150µl Triton-X buffer (1% Triton X-100, 150mM NaCl, 20mM HEPES pH 7.4, 1mM EDTA, 1.5mM MgCl_2_, 10% glycerol, 1X protease inhibitor cocktail, 50mM NaF, 2mM Na_3_VO_4_, 0.5mM PMSF) using a 200µl pipette followed by insulin syringes to mechanically break the hMOs. Lysates were incubated on an end-over-end rotator at 4°C for 30-40 minutes before being ultracentrifuged at 100,000 x g for 30 minutes at 4°C. Following ultracentrifugation, the supernatant was collected and labelled as the soluble fraction. The pellet was further resuspended in 150ul SDS buffer (2% SDS, 50mM Tris pH 7.4, 1X protease inhibitor cocktail) and boiled at 100°C for 10 minutes. The boiled samples were then sonicated using a Bioruptor Plus water bath sonicator set at low power for 10 cycles of 30 seconds ON and 30 seconds OFF. The sonicated samples were boiled again at 100°C for 10 minutes and ultracentrifuges at 100,000 x g for 30 minutes at room temperature. The supernatant was collected and labelled as the insoluble fraction. All protein concentrations were quantified using the DC Protein Assay (BioRad).

### Preparation of whole cell lysates from hMOs

Whole cell lysates were prepared by pooling 3 hMOs per condition followed by dissociation with a 200µl pipette and insulin syringes in RIPA buffer (ddH20, 50 mM Tris 7.4 pH, 150 mM NaCl, 1% NP40, 1X protease inhibitor cocktail). Lysates were incubated on an end-over-end rotator for 45 minutes at 4°C followed by centrifugation at 10,000 rpm for 10 minutes at 4°C. Supernatant was collected and protein concentration were quantified using the DC Protein Assay (BioRad).

### Immunoblot analysis

Lysates were mixed with 6X Laemmli buffer and boiled for 5 minutes, 10µg and 7µg protein was loaded in 12% and 10% polyacrylamide gels for α-syn and TH immunoblots respectively. SDS-PAGE separation was performed and proteins were transferred onto nitrocellulose membranes. To increase detection of α-syn, blots were fixed in 4%PFA + 0.1% glutaraldehyde for 30 minutes prior to blocking. Membranes were then blocked in a 5% milk solution in PBS buffer containing 0.2% Tween-20 (PBS-Tween) for 1 hour at room temperature, if membranes were fixed with PFA they were first washed 3 times in PBS-Tween prior to blocking. The membranes were then incubated in primary antibodies (**Supplementary table 2**) diluted in PBS-Tween overnight at 4°C. The next day the membranes were washed 3 x 5 minutes in PBS-Tween before incubation in peroxidase conjugated secondary antibody for 1 hour at room temperature. Membranes were again washed 3 x 5 minutes in PBS-Tween before image acquisition on the ChemiDoc MP System. Staining was then repeated for loading controls.

### Multi-electrode array (MEA) recordings

MEA plates (Axion Biosystems, CytoView MEA 24, M384-tMEA-24W) were coated with 10µg/ml Poly-L-Ornithine at 37°C overnight, followed by 5 µg/ml laminin at 37°C for 3 hours. 1 hour prior to recording, hMOs were placed on top of the electrodes with 20 µl of media to keep them from drying out. MEA plates were then put in the incubator for 1 hour to allow adequate attachment of hMOs on electrodes. After incubation, MEA plates were recorded for 10 minutes at a 12.5-kHz sampling frequency with an applied digital Butterworth bandpass filter. During recordings neurons were kept at 37 °C at 5% CO_2_. Electrode spikes were detected as any recorded spike that had a voltage of at least 6 SD away from background noise. All exported data were analyzed, and raster plots were generated using custom scripts written in Python (https://github.com/ghislainedeyab/AXION_spike_analysis) and the Axion Biosystems Neural Metric Tool. For pharmacological tests, all drugs used were diluted in the media of hMOs on the MEA and incubated for 15 minutes prior to recording again. For long term tetrabenazine treatment, tetrabenazine was added directly to hMO media in 6 well plates with every media change. All drugs used are listed in **supplementary table 3.**

### Flow Cytometry

Ten hMOs were gathered and combined into one gentleMACS M-Tube. hMOs were washed 3 times with 5 ml Dulbecco’s PBS (D-PBS). The D-PBS was removed and 2 mL of pre-warmed (37°C ) TrypLE express without phenol red was added to the tube and the tubes were placed on the automated GentleMACS Octo Heated dissociation device set at 37°C (24 minutes spin at 20rpm followed by 1 minute spin at 197rpm). After dissociation, 8 ml D-PBS was added to stop the enzymatic reaction and the cells were transferred to a 15 ml falcon tube through a 30 µm filter to get rid of clumps using low binding pipette tips. To capture leftover cells in the M-Tubes, 5 ml D-PBS was added into the M-Tubes and both sets of tubes were spun down for 5 minutes x 400 g. The supernatant was discarded and remaining cells in the M-Tube was resuspended with 1 ml D-PBS and was resuspended with the cells in the 15 ml falcon tube. All the cells were washed once more and resuspended in 1 ml D-PBS. The total cell number was counted using the Attune NxT Flow Cytometer (ThermoFisher). One million cells were recovered and resuspended in 1 ml D-PBS with 1 µl Live/Dead Fixable Dye for 30 minutes in the dark to assess cell viability. All following incubations were done in the dark to avoid loss of fluorescent signal. The single cell solution was then washed twice with D-PBS for 5 minutes x 400 g and resuspended with 5 µl Human TruStain FcX to block unspecific Fc receptor binding. The cells were washed once more and resuspended in FACS buffer (5% FBS, 0.1% NaN_3_ in D-PBS) and incubated for 30 minutes with fluorescent conjugated antibodies diluted in FACS buffer (**Supplementary Table 2**). The single cell solution was washed twice and fixed and permeabilized with 2% PFA in FACS buffer for 30 minutes followed by 0.7% Tween-20 in FACS buffer for 15 minutes to allow internalization of TH antibody. Following permeabilization, the cells were washed once more and incubated for 30 minutes with the fluorescent tagged TH antibody (**Supplementary table 2**). Cells were washed two more times and data acquisition was performed through the Attune NxT Flow Cytometer. All data were analyzed, and figures were produced using the CellTypeR analysis pipeline^37^ (https://github.com/RhalenaThomas/CelltypeR). A cell type marker expression matrix was generated specifically matching the antibody panel used in these experiments to include TH. Predictions of cell types for annotation were performed using the custom prediction matrix for the correlation prediction model and expression values of the markers overlapping with the CellTypeR marker panel for random forest and Seurat label transfer predictions.

### Statistical Analysis

All statistical analysis was performed using GraphPad Prism version 10.4.1 for MacOS, GraphPad Software, Boston, Massachusetts USA, www.graphpad.com. All analysis includes data from 3-4 independent hMO batches (N = batches) from each genotype. For MEA recordings 24 hMOs were used per batch, per genotype, and per timepoint (N = 4). For pharmacological testing, 6-8 hMOs were used per batch and per genotype (N = 3). For immunofluorescence and western blot analysis, 3-4 hMOs were used per batch and per genotype (N = 3). The chronic tetrabenazine treatment experiment was done on one batch of Iso CTL hMOs with 6 treated and 6 untreated hMOs. For Syn and pSyn immunofluorescence comparisons, analysis was done using a One-Way ANOVA followed by Tukeys multiple comparison test. For TH immunofluorescence comparisons, analysis was done using an unpaired T-test (two-tailed). Western blot analysis was done by normalizing α-syn or TH protein levels to respective Actin protein levels, followed by subsequent normalization of *SNCA* Trip hMOs to Iso CTL hMOs from the same batch, statistical analysis was done using a non-parametric Mann-Whitney test (one-tailed). For MEA data, outliers were removed using the ROUT method for outlier detection (Q = 1%), and the data was subsequently analyzed using a One-Way ANOVA with Tukeys multiple comparison test for time course analysis within the Iso CTL hMOs (**Fig.2B**), a Two-Way ANOVA with Sidaks Multiple Comparison test for time course experiments between genotypes (**Fig.3A**), or an unpaired T-Test (two-tailed) for bursting parameter comparisons between genotypes or separate batches of organoids (**Fig.3B-D, G-H**). A paired T-test (two-tailed) was used for comparison of hMO activity before and after drug treatment (**Fig.2D**)(**Fig.4A-D**) or unpaired T-test (two-tailed) for comparisons between genotypes (**Fig.4A-D**). All the data is presented as mean ± SEM.

## Data Availability

The datasets generated and analysed during the current study are available in the GitHub repository: https://github.com/ghislainedeyab/AXION_spike_analysis

## Acknowledgments

We would like to thank Xiuqing Chen for helping with iPSC quality control and facilitating access to the iPSC lines. We also want to thank The Neuro Microscopy Core Facility for confocal microscope management, maintenance, and training. We thank Meghna Mathur, Paula Lepine, and Nguyen-Vi Mohamed for their guidance on hMO generation and use in experimental assays. We would also like to acknowledge Eddie Cai for his help in writing and testing the macros used for IF analysis. This work was supported by a CIHR project grant (PJT-195804), a *Fonds d’Accéleration des Collaborations en Santé* grant from CQDM/MEI (Fon-FACS-013) and a Michael J. Fox Foundation grant (MJFF-021629). We gratefully acknowledge the funding support for this research from Parkinson Canada. G.D. received funding through McGill Healthy Brains for Healthy Lives (HBHL) graduate student fellowship, McGill Jeanne Timmins Costello Studentship, and the Parkinson Canada Graduate Student Award. TMD was supported by the Canadian New Frontiers in Research-Transformation (NRF-T) funded TRanslational Initiative to DE-risk NeuroTherapeutics (TRIDENT). E.A.F. is supported by a Canada Research Chair (Tier 1) in Parkinson’s disease. The funders were not involved in study design, data collection and analysis, or the writing of this manuscript.

## Author Contributions

The project was conceived by G.D. and E.A.F. G.D. cultured iPSCs, generated and cultured hMOs, collected and processed hMOs for immunofluorescence and immunoblot analysis, processed hMOs for flow cytometry, recorded hMOs on the MEA, and analyzed MEA data.

R.A.T. processed and analyzed flow cytometry data and provided guidance on custom scripts used for MEA data extraction and visualization. X.E.D. and G.D. created and tested custom scripts used for MEA data extraction and visualization. J.L. assisted in immunofluorescence assays and confocal imaging. J.S. provided guidance and technical assistance in flow cytometry experiments. Z.A. and N.S. provided organoids used for soluble/insoluble protein fractionation and synuclein western blot. Data were interpreted by G.D., R.A.T. and E.A.F. The manuscript was written by G.D. and E.A.F. with contributions by R.A.T., T.M.D., and Z.A. The project was supervised by E.A.F.

