## Supplementary Tables for "Midbrain organoids with an *SNCA* gene triplication display dopamine-dependent alterations in network activity"

**Supplementary Table 1**

| Reagent | Company | Cat Number |
|---|---|---|
| 2-mercaptoethanol | Merck | 8057400005 |
| Accutase | StemCell | 07922 |
| Antibiotic-Antimycotic | Gibco | 15240-062 |
| Aqua-poly/mount | PolySciences | 18606-20 |
| Ascorbic acid | Sigma | A5960 |
| B27 without vitamin A | Gibco | 12587010 |
| BDNF | Peprotech | 450-02 |
| Bioreactors | Corning | 4500-500 |
| CHIR99021 | Selleckchem | S2924 |
| Db-cAMP | Carbosynth | ND07996 |
| DMEM/F12 | Gibco | 10565-018 |
| Dulbeccos PBS | Multi-cell | 311-425-CL |
| EB Disc 360 well ULA-Coated | eNUVIO | eN-eb360u-001 |
| FGF-8 | Peprotech | 100-25 |
| GDNF | Peprotech | 450-10 |
| Glutamax | Gibco | 35050-061 |
| Heparin | Sigma | H3149 |
| Insulin | Sigma | I2643 |
| Laminin (media) | Sigma | L2020 |
| Laminin (coating) | Invitrogen | 23017-015 |
| Matrigel | Corning | 354277 |
| Matrigel reduced growth factor | Corning | 356230 |
| MEM-NEAA | Wisent | 321-011-EL |
| mTeSR 5X Supplement | StemCell | #85850 (component #85852) |
| mTeSR Basal media | StemCell | #85850 (component #85851) |
| N2 | Gibco | 17502048 |
| Neurobasal | Life Technologies | 21103-049 |
| Noggin | Peprotech | 120-10C |
| OCT tissue embedding compound | Thermo Fisher | 23730571 |
| Penicillin-Streptomycin | Wisent | 450-200-EL |
| Paraformaldehyde | Thermo Fisher | 28908 |
| Poly-L-Ornithine | Sigma | P3655 |
| Protease inhibitor cocktail | Millipore Sigma | 11836170001 |
| SB431542 | Selleckchem | S1067 |
| SHH | Peprotech | 100-45 |
| Triton X-100 | Fisher Scientific | BP151-500 |
| TrypLE | Gibco | 12604-013 |
| Tween-20 | Thermo Fisher | BP337-500 |
| Ultra low attachment 6-well plates | Corning | 3471 |
| Y-27632 | Selleckchem | S1049 |

**Supplementary Table 2**

| Antibody | Company | Cat Number | Application | Concentration |
|---|---|---|---|---|
| TH anti-Rb | Pel-Freez | P40101 | IF/WB | 1:500 |
| Actin anti-Ms | Millipore | MAB1501 | WB | 1:50,000 |
| Syn anti-Ms | BD Bioscience | 610787 | IF | 1:500 |
| pSyn EPI anti-Rb | Abcam | ab51253 | IF | 1:500 |
| Syn MJFR anti-Rb | Abcam | ab138501 | WB | 1:1000 |
| MAP2 anti-Ch | Encor Biotech | CPCA-MAP2 | IF | 1:1000 |
| Hoechst | Invitrogen | H3570 | IF | 1:5000 |
| GFAP anti-Rb | DAKO | Z033429-2 | IF | 1:500 |
| Nestin anti-Ms | Invitrogen | Ma1-110 | IF | 1:500 |
| GAD67 anti-Ms | Millipore | MAB5406 | IF | 1:500 |
| CD140a | R&D Systems | FAB1264N | Flow Cytometry | 1:320 |
| CD44 | Biolegend | 338810 | Flow Cytometry | 1:320 |
| SSEA4 | Biolegend | 330418 | Flow Cytometry | 1:80 |
| CD133 | Biolegend | 372810 | Flow Cytometry | 1:320 |
| CD184 | Biolegend | 306522 | Flow Cytometry | 1:640 |
| CD15 | Biolegend | 323044 | Flow Cytometry | 1:80 |
| CD29 | Biolegend | 303014 | Flow Cytometry | 1:160 |
| CD56 | Biolegend | 392420 | Flow Cytometry | 1:320 |
| CD24 | Biolegend | 311136 | Flow Cytometry | 1:160 |
| TH | Invitrogen | MA5-38641 | Flow Cytometry | 1:1780 |
| Live/Dead Fixable Dye | Invitrogen | L34966 | Flow Cytometry | 1:2000 |
| Human TruStain FcX | Invitrogen | 422303 | Flow Cytometry | 1:20 |

**Supplementary Table 3**

| Drug | Company | Cat Number |
|---|---|---|
| Tetrodotoxin | Sigma | T5651 |
| Quinpirole | Tocris | 1061/10 |
| Tetrabenazine | Sigma | T2952 |
